# Sexually dimorphic dynamics of the microtubule network in medaka (*Oryzias latipes*) germ cells

**DOI:** 10.1101/2023.04.09.536194

**Authors:** Mariko Kikuchi, Miyo Yoshimoto, Tokiro Ishikawa, Yuto Kanda, Kazutoshi Mori, Toshiya Nishimura, Minoru Tanaka

## Abstract

Oogenesis is the process through which ovarian germ cells enter meiosis and acquire essential characteristics of an egg, such as an increase in follicular size and deposition of maternal factors. In vertebrates, however, the pathways that initiate oogenic processes after germ cell sex determination remain elusive. Here we report a dome-shaped microtubule structure that is maintained during oogenesis. This structure is established in primordial germ cells and maintained in gonocytes and stem-type germ cells in a *foxl3*-independent manner. *dmrt1* is required for disruption of the dome structure in differentiating spermatogonia. In addition, the dome structure accumulates mitochondria and Golgi apparatus, and is replaced by the Balbiani body, an oocyte-specific aggregate producing animal-vegetal polarity, suggesting a role for this structure in organelle transfer and oocyte polarization. Our findings show the presence of a pathway regulating a sexually dimorphic microtubule network independent of germ cell feminization by *foxl3*.

**Summary Statement:** A dome-shaped microtubule structure leading to oocyte polarity was identified in germ cells. Genetic pathways involved in the formation and destabilization of the structure during germ cell sex determination were clarified.

## Introduction

Germ cells differentiate into dimorphic gametes, eggs and sperm, and these are the only cells that transmit genetic information to the next generation. Therefore, the mechanism of gametogenesis has been intensively studied. After the commitment of germ cells to oogenesis (female sex determination of germ cells), several pathways need to be triggered to implement the essential characteristics for functional eggs. These include increasing cellular size and depositing maternal factors for embryogenesis. In vertebrates, however, the initial pathways that are integrated and triggered at the time of germ cell sex determination are largely not known.

Unlike mammals, the existence of sexually indifferent germline stem cells has been demonstrated in female teleost fish, medaka (*Oryzias latipes*) (Nakamura et al., 2010), and Foxl3 (Forkhead box L3), a germ cell intrinsic factor that determines oogenesis or spermatogenesis was identified (Nishimura et al., 2015). In addition, the sex of the medaka is genetically determined by the presence or absence of the sex determination gene, *DMY*/*dmrt1bY* on the Y chromosome (Matsuda et al., 2002; Nanda et al., 2002). Medaka are therefore suitable for analyzing the mechanism of germ cell sex determination and early gametogenesis (Nishimura and Tanaka, 2016; Kikuchi and Tanaka, 2022); two pathways of oogenesis triggered by *foxl3* have been identified. *foxl3* activates *fbxo47* and *rec8a* independently (Kikuchi et al., 2019; Kikuchi et al., 2020) and a genetic pathway downstream of *fbxo47* leads to the formation of follicles while *rec8a* is essential for the progression of meiosis only during oogenesis.

Another important pathway for oogenesis is the creation of oocyte polarity. In non-mammalian vertebrates, oocyte polarity lays the foundation of the embryonic axis. In *Xenopus* and zebrafish, the animal-vegetal axis of the egg is established in early oocytes via a structure called the Balbiani body (Bb) (Elkouby, 2017; Jamieson-Lucy and Mullins, 2019). The Bb localizes mRNAs encoding dorsal determinants and related factors vegetally in oocytes (Nojima et al., 2010; Lu et al., 2011; Ge et al., 2014). The lack of a Bb structural component, Bucky ball (Buc), results in symmetrical eggs that fail to develop beyond the early cleavage stages (Marlow and Mullins, 2008). These results demonstrate that Bb formation is an essential process for functional egg development (Elkouby et al., 2016).

In this study, we identified a dome-shaped microtubule (MT dome) structure that is formed in primordial germ cells (PGCs) and is maintained in stem-type germ cells in both sexes. The structure is destabilized during spermatogenesis while it persists during oogenesis and is associated with Bb formation at the zygotene stage. Interestingly, our genetic analysis indicates that the structure is not under the control of *foxl3* during oogenesis, representing a pathway that potentially contributes to egg polarity. In addition, we describe the genes involved in the maintenance and destabilization of the MT dome structure.

## Results

### Regulation of the stabilization of the MT dome differs between oogenesis and spermatogenesis

Our previous transcriptome analysis revealed that the expression of genes related to the microtubule regulatory pathway changes significantly downstream of *foxl3* (Kikuchi et al., 2019). This led us to hypothesize that the organization of the microtubule network in germ cells is different between the sexes during gametogenesis. To explore the microtubule network in germ cells, we labeled microtubules with antibodies against pan and acetylated α-tubulins. Both antibodies detected a conspicuous dome-shaped structure localized at the perinuclear cytoplasmic region in oogonia and spermatogonia at the hatching stage (**Fig. S1**). Construction of a 3D image with co-staining of γ-tubulin revealed that the MT dome is hollow and forms around the centrosome(s), which was always located at the base of the MT dome (**Movie 1; Fig. S2**).

We then tracked the structure of the MT dome during gametogenesis. During oogenesis, a single stem-type germ cell (type-I oogonia) differentiated to form an oogonial cyst composed of 8–32 cells (type-II oogonia) after 3–5 rounds of mitosis with incomplete cytokinesis (Saito et al., 2007; Nakamura et al., 2010). The MT dome was consecutively observed from type-I oogonia to meiocytes at the zygotene stage (**Fig. 1A-D; Movies 2-4**). Interestingly, the MT domes in mitotically dividing type-II germ cells in a single cyst were connected by a bundle of acetylated microtubules via intercellular bridges (**Fig. 1B; Movie 2**). This connection was kept until the leptotene stage, and lost by the zygotene stage (**Fig. 2C-D; Movies 3-4**). From the pachytene stage onward, acetylated microtubules were arranged into a cage-like structure that surrounds the nucleus (**Fig. 1E; Movies 5-6**).

**Figure 1.**
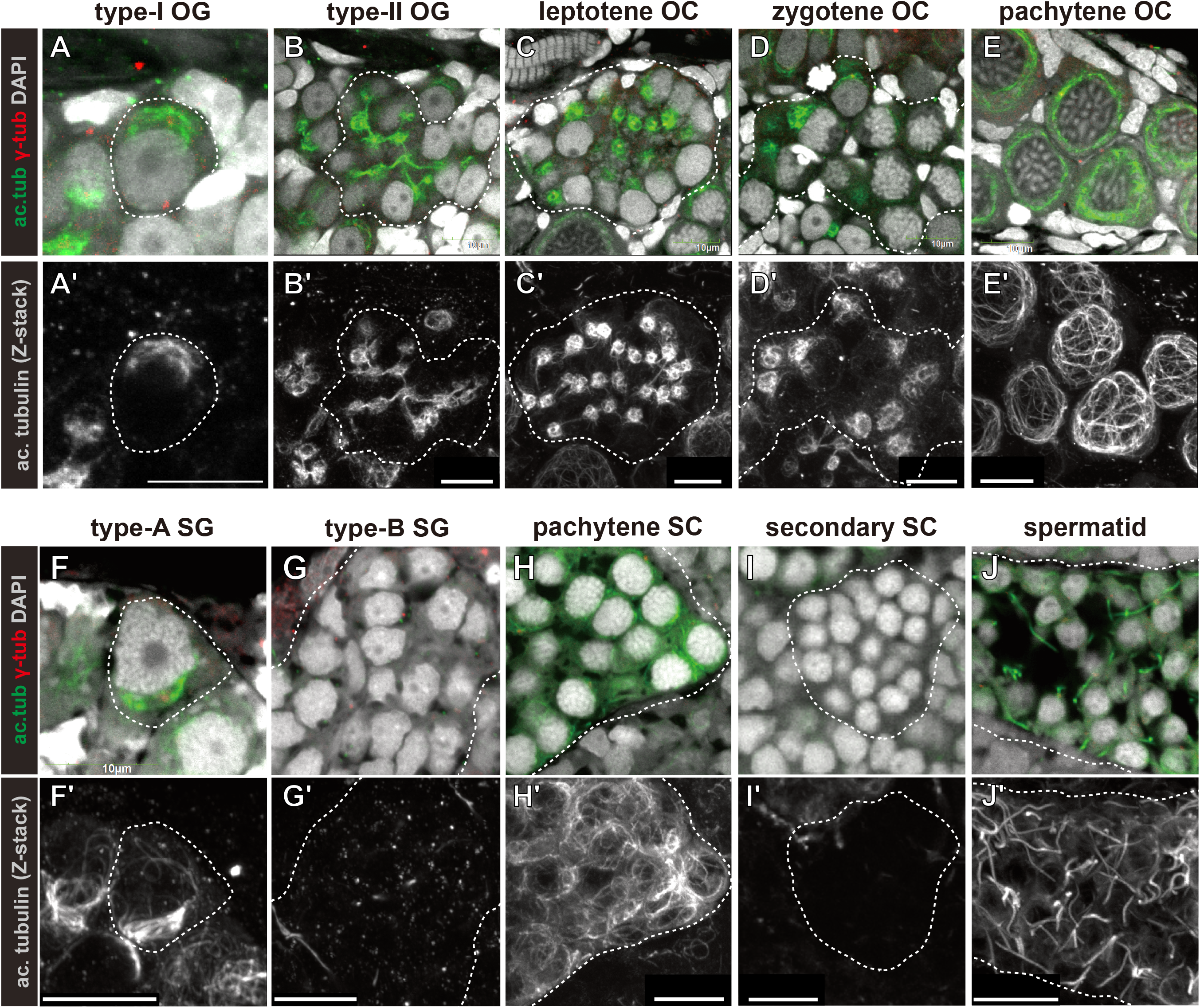
Sexually dimorphic dynamics of the microtubule network during oogenesis and spermatogenesis. (A-J) Immunofluorescence of 10 dph XX ovaries (A-E) and adult XY testes (F-J). Dotted lines indicate single germ cells (A and F) or single cysts of germ cells (B-D, G-J). Bottom panels show z-stacked gray-scale images of acetylated α-tubulin. The MT dome was observed in type-I oogonia (OG) (A) and type-A spermatogonia (SG) (F). The structure was maintained in type-II OG and oocytes (OC) until the zygotene stage (B-D), while it was destabilized in type-B SG (G). In pachytene OC, microtubules were organized into a cage-like structure encompassing the nucleus (E). The MT dome was not observed in primary spermatocytes (SC) (pachytene) (H), secondary SC (I), and spermatids (J). Acetylated α-tubulin signals in J show forming sperm flagella. Scale bars: 10 µm.

**Figure 2.**
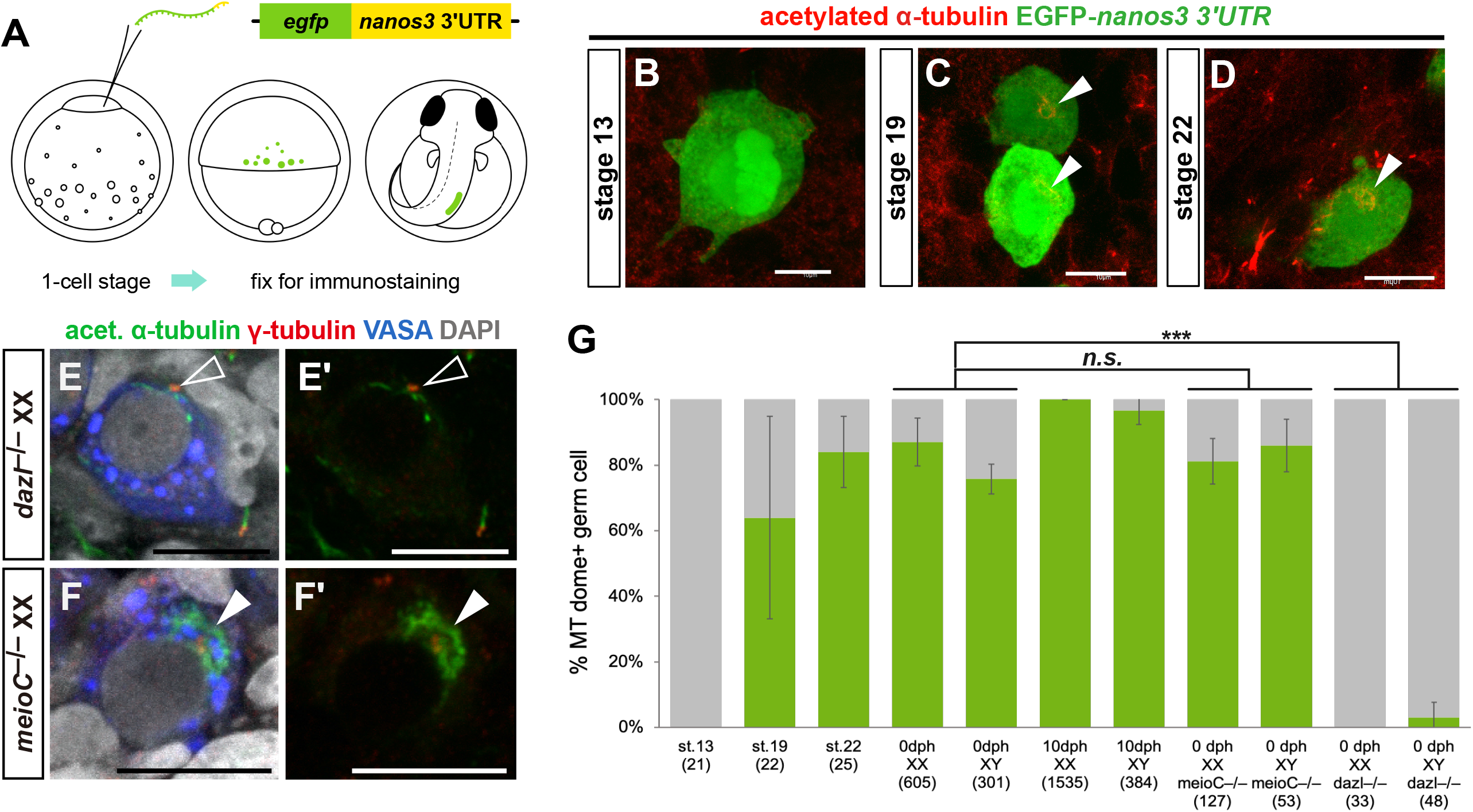
The MT dome is formed in primordial germ cells and is lost in gonocytes of *dazl* mutants. (A) PGCs were labeled with EGFP by injecting *egfp*-*nanos3* 3’UTR mRNA into 1-cell-stage embryos. (B–D) Immunohistochemistry of embryos at stage 13 (B; early gastrula stage), stage 19 (C; 2-somite stage), and stage 22 (D; 9-somite stage) for acetylated α-tubulin and EGFP. Arrowheads indicate the MT domes. Scale bars: 10 µm. (E and F) The MT dome was not observed in *dazl*^−/−^ XX germ cells (E and E’, open arrowheads) but was retained in *meioC*^−/−^ XX germ cells (F and F’, filled arrowheads). Scale bars: 10 µm. (G) Quantification of the number of germ cells containing the MT dome in developing gonads of the wild-type and mutants. Since gonadal sex differentiation begins at stage 33, the genetic sex of stage 13–22 embryos was not specified. For 0–10 dph XX samples, only mitotic germ cells were counted. Numbers in brackets indicate the number of germ cells quantified from four to six gonads. Data are shown as mean ± SD. *** *p* < 0.001.

During spermatogenesis, a single stem-type germ cell (type-A spermatogonia) mitotically proliferated to form a cyst containing 128–1024 differentiating spermatogonia (type-B spermatogonia) (Shibata and Hamaguchi, 1988; Sumita et al., 2022). As in females, the MT dome was detected in type-A spermatogonia (**Fig. 1F**). However, it was destabilized in type-B spermatogonia (**Fig. 1G**). When type-B spermatogonia entered meiosis, acetylated microtubules transiently reappeared in primary spermatocytes, but were lost in secondary spermatocytes (**Fig. 2H-I**). Of note, the microtubule structure in the primary spermatocytes was not like that of the MT dome; rather it was more obscure and distributed broadly in the cytoplasm. It has been reported that the microtubule network is required for chromosomal arrangement during meiotic prophase I, which facilitates synapsis between homologous chromosomes (Scherthan, 2001; Chikashige et al., 2006; Shibuya et al., 2014). Thus, the transient increase of microtubules in medaka primary spermatocytes may be related to progression of meiosis. During spermiogenesis, acetylated microtubules were strongly detected on the forming flagella (**Fig. 2J**). Collectively, although the MT dome is detected in stem-type germ cells of both sexes, it is maintained only in female germ cells from the differentiating stage to early meiotic prophase I. These observations indicate that regulation of the stabilization of the MT dome differs according to sex.

### The MT dome is initially formed in PGCs and maintained in gonocytes and stem-type germ cells

Next, we analyzed the developmental time course of MT dome formation. PGCs were labeled with EGFP by injecting synthetic *egfp* mRNA fused with *nanos3* 3’UTR (*egfp*-*nanos3* 3’UTR) into 1-cell-stage embryos (**Fig. 2A**, Kurokawa et al., 2006; Kikuchi et al., 2020). These embryos were used for immunostaining with antibodies for EGFP and acetylated tubulin. At embryonic stage 13 (early gastrulation stage), when germ cell lineage is established (Kurokawa et al., 2006), no MT dome was observed in early PGCs (**Fig. 2B, G**). At stage 19 (2-somite stages), when PGCs are posteriorly migrating along the lateral plate mesoderm (Nakamura et al., 2006), the MT dome was first observed in approximately 64% of PGCs (**Fig. 2C, G**). PGCs containing the MT dome increased to 84% by stage 22 (9-somite stage), and it was consistently observed in gonocytes at the timing of gonadal formation (stage 30–33) and gonadal sex determination (stage 33–35) (**Fig. 2D, G, Fig. S3**). By 10 days post-hatching (dph) when female germ cells enter meiosis while male germ cells continue mitotic cell divisions (Nishimura and Tanaka, 2014), most XX and XY mitotic germ cells, including stem-type germ cells, had the MT dome (**Fig. 2G; Fig. S4**). Collectively, the MT dome first appears in PGCs by stage 19, and is maintained in both XX and XY gonocytes and stem-type germ cells.

### The MT dome in PGCs and gonocytes is maintained in a dazl-dependent manner

In some vertebrates, including mammals and fishes, *dazl* (*deleted in azoospermia like*) and *meioC* (*meiosis specific with coiled-coil domain*) are involved in gonocyte formation and gametogenesis commitment, respectively (Gill et al., 2011; Abby et al., 2016; Soh et al., 2017; Nishimura et al., 2018). PGCs in *dazl* mutants do not develop normal gonocytes. In the absence of *meioC* function, stem-type germ cells accumulate and no meiotic germ cells are observed. We investigated if *dazl* or *meioC* are involved in the persistence of the MT dome using *dazl*^−/−^ and *meioC*^−/−^ gonads (Nishimura et al., 2018). The number of germ cells containing the MT dome significantly decreased in *dazl*^−/−^ XX and XY gonads, while there was no significant reduction of germ cells with the MT dome in *meioC*^−/−^ compared to the wild type (**Fig. 2E-G**). These results demonstrate that *dazl,* but not *meioC,* is required for the maintenance of the MT dome in the gonad.

### Stability of the MT dome is regulated downstream of dmrt1 but not foxl3

In medaka, the sexual fate of germ cells is determined by *foxl3*. Loss of function of *foxl3* leads to spermatogenesis in XX ovaries (Nishimura et al., 2015). To investigate whether *foxl3* regulates the stabilization of the MT dome in differentiating type-II oogonia, *foxl3*^−/−^ XX ovaries were immunostained with acetylated tubulin. Similar to wild-type XX, both MT domes and their connections by microtubules within a cyst were observed normally in the spermatogenesis-committed germ cells of *foxl3*^−/−^ XX mutants (**Fig. 3A, B**). This indicates that *foxl3* is not required for maintenance of the MT dome. In addition, the mutants showed that the presence of the MT dome does not impede progression of spermatogenesis occurring in *foxl3*^−/−^ ovaries (**fig. 3B**) (Nishimura et al., 2015).

**Figure 3.**
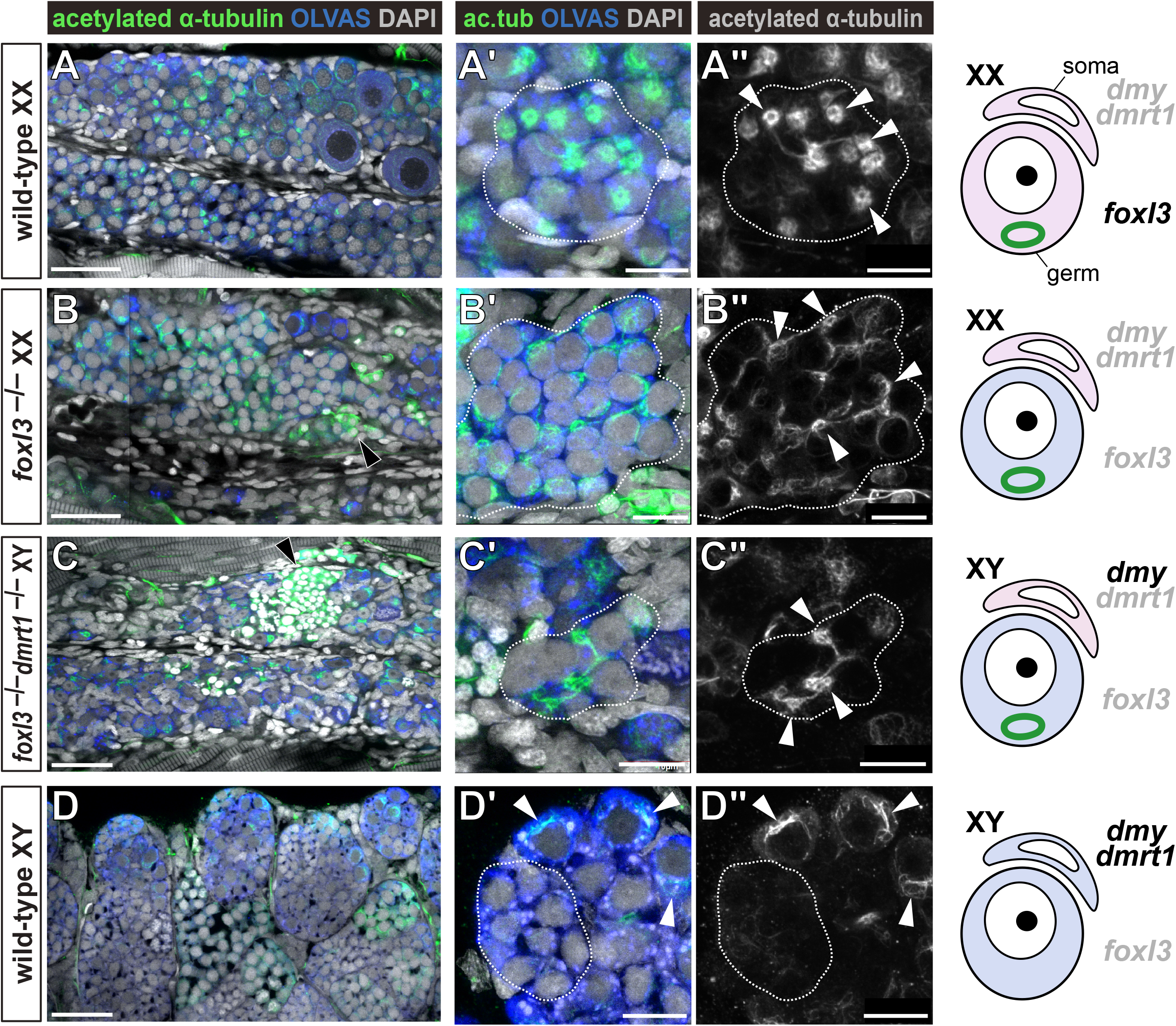
Destabilization of the MT dome in spermatogonia is regulated downstream of *dmrt1*. (A-D) Immunofluorescence of 10 dph gonads of wild-type XX (A-A″), *foxl3*^−/−^ XX (B-B″), *foxl3*^−/−^, *dmrt1*^−/−^ XY (C-C″), and wild-type XY (D-D″). The middle two columns are z-stack images of a single cyst indicated by dotted lines. The right column illustrates the sexual fate of somatic cells (soma) and germ cells (germ). Pink and blue colors indicate female- and male-fated cells, respectively. In XX (A and B), the MT dome was maintained independently of a *foxl3*-pathway. In *foxl3*/*dmrt1* double mutant XY (C), the MT dome was observed in masculinized germ cells, while it was destabilized in wild-type type-B spermatogonia (D). Black arrowheads in B and C indicate spermatids. White arrowheads in D’ and D″ indicate the MT domes in type-A spermatogonia. Scale bars: 30 µm (A-D) or 10 µm (A’-D’, A″-D″)

In wild-type testes, *dmrt1* is expressed in gonadal somatic cells and required for masculinization of gonads. Depletion of *dmrt1* causes upregulation of female somatic genes including *foxl2* and *aromatase*, resulting in male-to-female sex reversal of XY fish (Masuyama et al., 2012). We hypothesized that the *dmrt1* negatively regulates the MT dome in XY germ cells. To explore this possibility, the MT dome was examined in *dmrt1*/*foxl3*-double knockout (dKO) XY gonads, in which *dmrt1*^−/−^ somatic cells become feminized while *foxl3*^−/−^ germ cells intrinsically commit to spermatogenesis. We found that the MT dome was stabilized in type-B spermatogonia in *dmrt1*/*foxl3*-dKO XY gonads (**Fig. 3C**). This indicates that the *dmrt1*-downstream pathway promotes destabilization of the MT dome in wild-type XY germ cells (**Fig. 3D**).

### The MT dome enriched with organelles is replaced with a Balbiani body

The Bb is an oocyte-specific non-membranous aggregate containing mRNA, proteins, and organelles (*e.g.*, mitochondria, ER, Golgi apparatus) (Pepling et al., 2007). In *Drosophila*, zebrafish, and mice, microtubule-mediated organelle transport among differentiating cyst cells is essential for Bb formation, which promotes oocyte development (Roper and Brown, 2004; Elkouby et al., 2016; Lei and Spradling, 2016; Niu and Spradling, 2022). We labeled mitochondria with a CFP-tag fused with a mitochondrial targeting signal (CFP-mito) and found that the CFP-mito signal was enriched around the MT dome as well as the germinal granule in stem-type type-I and differentiating type-II oogonia (**Fig. 4A**). From zygotene to diplotene oocytes, the mitochondrial signal dispersed and localized to a cage-like microtubule structure around the nucleus (**Fig. 4A**). Interestingly, the timing of the loss of mitochondrial accumulation is consistent with that of loss of the microtubule extension that connects the MT domes (**Fig. 1D**). In transmission electron micrographs (TEM), the Golgi apparatus was also localized in the area of the MT dome with a circular disposition around centrosomes in the oogonia (**Fig. 4B, Fig. S5**). These observations suggest that distribution of mitochondria and the Golgi apparatus is associated with the MT dome network.

**Figure 4.**
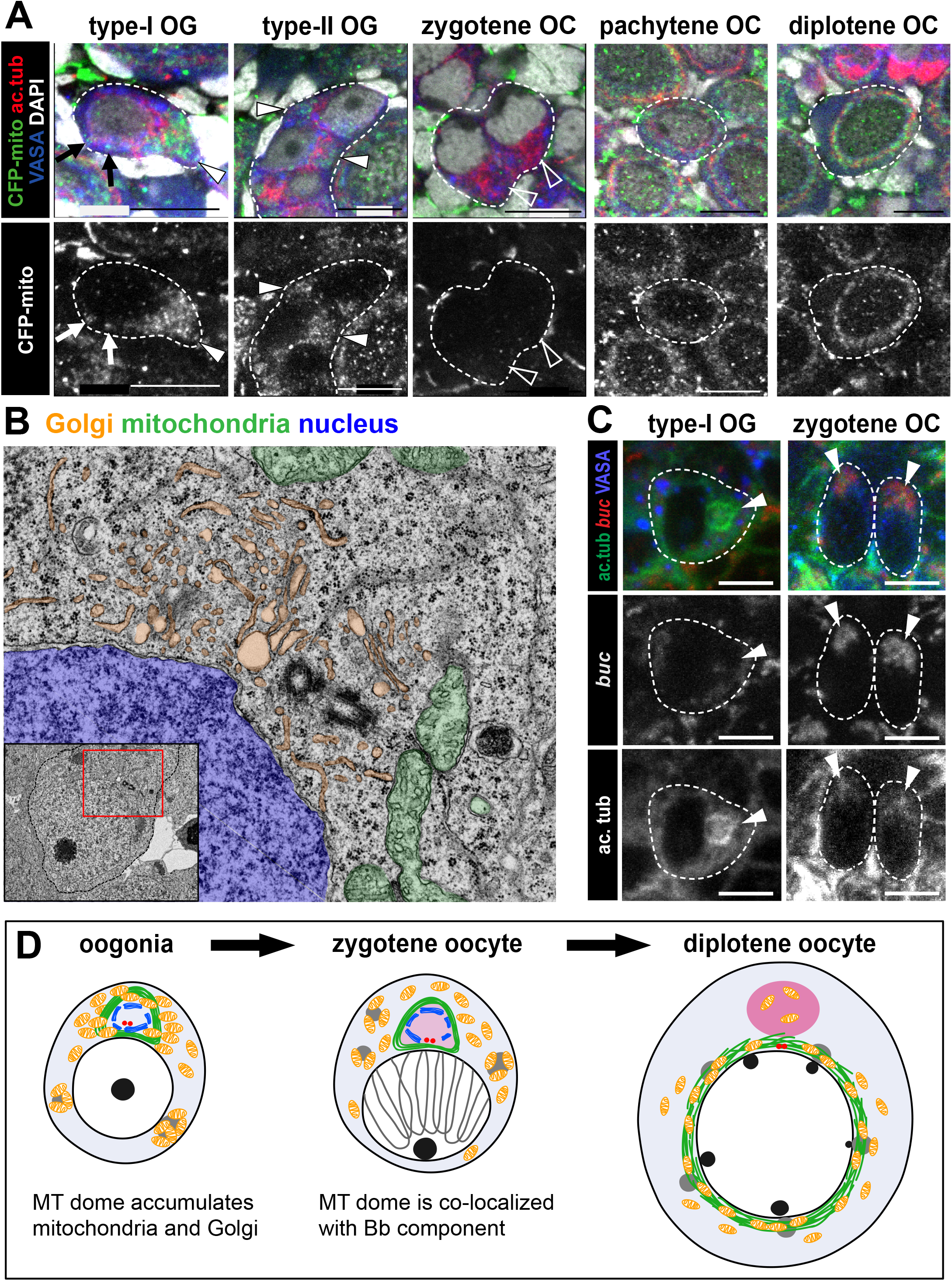
The MT dome is located in an organelle-rich region and is enriched with *buc* mRNA. (A) Localization of mitochondria (CFP-mito) around the MT domes (arrowheads) and germinal granules (arrows) in type-I and type-II oogonia. Mitochondrial accumulation was not observed in zygotene oocytes but was found in the MT cage in pachytene and diplotene oocytes. Scale bars: 10 µm. (B) TEM image of type-I oogonia at 10 dph. Cytoplasmic region indicated in the inset is magnified. Golgi apparatus, mitochondria, and nucleus are colored orange, green, and blue, respectively. Centrioles are located at the center. (C) Co-staining of acetylated α-tubulin and *buc* mRNA in type-I oogonia and zygotene oocytes. Arrowheads indicate the MT domes. Accumulation of *buc* mRNA was detected from the zygotene stage. Scale bars: 5 µm. (D) Graphical summary of mitochondria (orange), Golgi apparatus (blue), and *bucky ball* (pink) localization during oogenesis. In oogonia, mitochondria and Golgi accumulate around the MT dome (green) as well as germinal vesicles (gray). In zygotene oocytes, Golgi disposition is retained, while mitochondrial accumulation around the MT dome is lost. *buc* mRNA is co-localized with the MT dome at zygotene stage. By the diplotene stage, the MT dome is replaced with Bb, and microtubules are organized into a cage-like structure around nucleus.

Like the Bb, the MT dome is consistently present around centrosomes at the nuclear periphery (**Fig. S2**). In zebrafish and mice, polarized distribution of the Bb component along microtubules is initially observed at around the zygotene stage (Elkouby et al., 2016; Niu and Spradling, 2022). On the other hand, the MT dome in medaka was formed as early as the stage of migrating PGCs and was maintained in females at the zygotene stage (**Fig. 1A-E**). Based on these observations, we postulated that the MT dome may act as a precursor to Bb so that structural components of Bb can accumulate during meiosis I. Buc is known to be required for Bb formation and functions in Bb assembly as an intrinsically disordered protein comprising a lattice-like meshwork (Bontems et al., 2009; Elkouby, 2017). In medaka, Buc is encoded by the *buc* and *bucl* genes (**Figs. S6, 7**). Whole-gonad *in situ* hybridization revealed that *buc* mRNA is initially detected in zygotene oocytes and accumulates in the Bb in previtellogenic oocytes (**Fig. S8**). In addition, *buc* mRNA was co-localized with the MT dome in zygotene oocytes (**Fig. 4C**), suggesting that the MT dome we found frames the structure of the Bb. Collectively, these data support the idea that the MT dome promotes oocyte development via organelle transfer and Bb formation (**Fig. 4D**).

## Discussion

We revealed a unique and conspicuous shape of a microtubule structure in medaka germ cells, which is formed as early as the somitogenesis stage when PGCs are migrating toward gonads. 3D analysis revealed that its shape is dome-like and harbors a centrosome inside (MT dome) (**Fig. S2**). In mitotically dividing oogonia, microtubule bundles extend from the MT dome through an intercellular bridge to other MT domes of adjacent germ cells within a single cyst. Interestingly, the MT dome is abolished in type-B spermatogonia, suggesting its importance in oogenesis, and is present until it is replaced with Bb (**Fig. 5A**).

**Figure 5.**
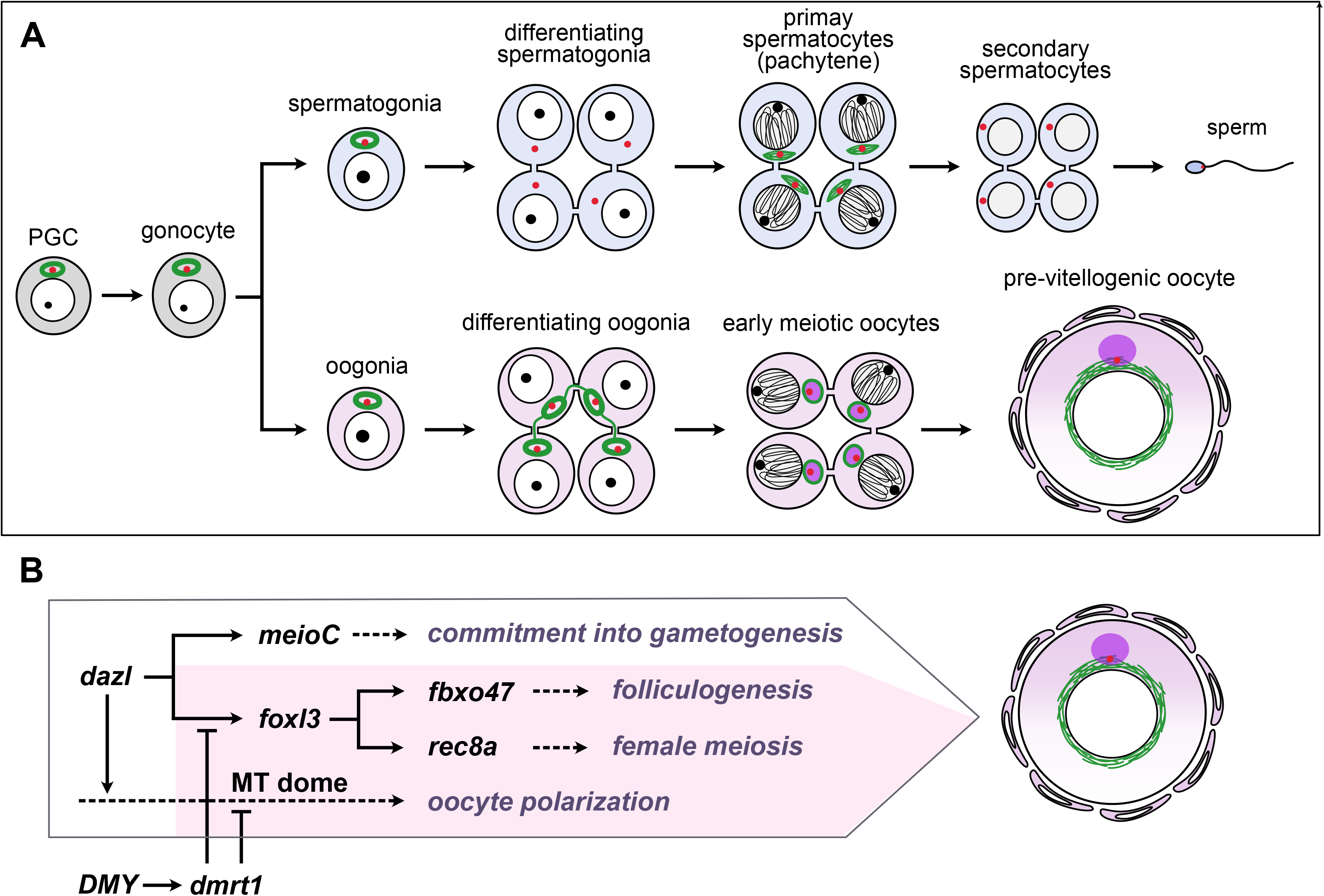
Graphical summaries of germline sexual development and genetic pathways regulating oogenesis in *Oryzias latipes*. (A) Sexually dimorphic changes to the microtubule network during gametogenesis. The MT dome is formed in PGCs and maintained in gonocytes and germline stem cells. During oogenesis, the MT dome persists until the zygotene stage, and is replaced with Bb components such as *bucky ball*. On the other hand, the MT dome is abolished in differentiating type-B spermatogonia. Acetylated α-tubulin (green), centrosome (red), and Bb (purple) are shown. (B) Genetic pathways regulating oogenesis. Each pathway contributes to creating a distinct feature of oogenesis, including commitment to gametogenesis, folliculogenesis, female meiosis, and oocyte polarization. In this study, we revealed a *foxl3*-independent pathway leading to oocyte polarity. *dazl* is required for the presence of the MT dome in gonocytes, while *dmrt1* promotes destabilization of the MT dome in male germ cells.

Although the requirement of the microtubule structure in oocyte development is conserved in animals including *Drosophila* (Cox and Spradling, 2003), zebrafish (Elkouby et al., 2016), and mice (Lei and Spradling, 2016; Niu and Spradling, 2022), genetic pathways that regulate dynamic changes of microtubule arrangements remain largely elusive in vertebrates.

Contrary to our initial expectation, persistence of the MT dome during oogenesis does not require *foxl3,* which is essential for germline stem cells to commit to oogenesis. Instead, we found that *dmrt1* contributes to the loss of the MT dome during spermatogenesis (**Fig. 5B**), indicating that the MT dome is actively degraded during germline masculinization. Meanwhile, in the absence of *dazl*, germ cells fail to retain the MT dome (**Fig. 2E**, Nishimura et al., 2018), suggesting the requirement of *dazl* function in germ cells that reached the gonad. Our data also suggests that *meioC* is not required for the MT dome formation in type-I germ cells (**Fig. 2F**), although the possibility remains in that *meioC* acts to maintain the MT dome in differentiating oogonia. Collectively, a pathway maintaining the MT dome, leading to female-specific Bb, is independent of *foxl3*. These observations suggest that germ cells implement oocyte characteristics before the sex of the germ cell is determined.

Based on previous reports and our data, two functions can be considered for the MT dome. First, we showed that the MT dome in medaka is positioned in an organelle-enriched area and extends MT bundles towards adjacent germ cells in a single cyst via intercellular bridges (**Fig. 1B-C**). This association disappears as intercellular connections are lost during meiotic prophase I (**Fig. 4A**). This is consistent with reports that some germline cyst cells transfer their cytoplasmic component to pro-oocytes via an intercellular bridge in a microtubule-dependent manner (Lei and Spradling, 2016; Niu and Spradling, 2022). It is possible that the MT dome in medaka provides a scaffold for the MT extension connecting different cystic germ cells and is consequently involved in organelle and cytoplasmic transfer.

Regarding this point, it is interesting to note here that the fertilization rate of *foxl3*^−/−^ sperm is not as high as that of the wild type (Nishimura et al., 2010). This may be due to the persistence of the MT structure in differentiating spermatogonia in *foxl3*^−/−^ ovaries, although the presence of the MT dome did not impede the progression of spermatogenesis (**Fig. 3B**). The presence of the MT dome may affect the quality of sperm.

Second, as discussed above, the structure may be a prerequisite for Bb formation because an organelle aggregates around the MT dome and, as the MT dome is rearranged with a cage-like structure surrounding the oocyte nucleus, Bb replaces the MT dome. In zebrafish and *Xenopus*, mRNAs encoding dorsal determinants, such as *wnt8a* and *grip2a*, are localized to Bb and contribute to generating the animal-vegetal axis (Marlow and Mullins, 2008; Lu et al., 2011; Ge et al., 2014). Therefore, it is likely that the MT dome predetermines the location and sets a frame for Bb as the MT dome is transformed into a cage-like structure surrounding the oocyte nucleus from the zygotene to pachytene stages.

## Materials and methods

### Fish

The OKcab strain of medaka fish (*Oryzias latipes*) was used in this study. Fishes were maintained in fresh water at 25-28°C under photoperiodically regulated conditions (14 hours light and 10 hours dark). Developmental stages of embryos were determined according to Iwamatsu (2004). All experiments were conducted with the approval of the Nagoya University official ethics committee (Approval Number S220008-002 in Department of Science, Nagoya University). Genotypic sex of all animals was determined by qPCR using TaqMan MGB probes (Thermo Fisher Scientific) that detect the male-determining gene *DMY*/*dmrt1bY* (Matsuda et al., 2002; Nanda et al., 2002) and the autosomal gene *cyp19a1* as described previously (Kikuchi et al., 2020). Generation and identification of the mutant alleles of *dazl* (ex2-Δ34) (Nishimura et al., 2018), *meioC* (ex5-Δ25) (Nishimura et al., 2018), and *foxl3* (NΔ17) (Nishimura et al., 2015) were described previously.

### Generation of dmrt (Δ13) mutant by TALEN-induced mutagenesis

The TALEN target sites for *dmrt1* exon 2 were searched using TALEN Targeter program (https://tale-nt.cac.cornell.edu/node/add/talen) with the following parameters: spacer length of 15-18 bp, repeat array of 16-18 bp, and upstream base of T only. TALEN assembly was performed following a modified version of the original protocol (Sakuma et al., 2013). *dmrt1*-target sites are as follows: right: TTCCGGAGGGCCCGGC, left: GTCCCCCCGGATGCCCA. TALEN plasmids were linearized by NotI digestion and used as templates for *in vitro* RNA synthesis with the mMESSAGE mMACHINE T7 transcription kit (Thermo Fisher Scientific). TALEN mRNAs (250 ng/μl left and right) were injected into one- or two-cell stage embryos. The F0 founders were crossed with wild-type medaka. A mutant allele with 13 bp deletion (deleted sequence: GGCTCCGGCTCCA; *dmrt1*Δ13) was identified in F1 adult fish using the primer sets shown in **Table S1** and sequence analysis. *dmrt1*Δ13^+/–^ fishes were crossed with *foxl3*Δ17^+/–^ fishes to obtain *foxl3*Δ17^+/–^; *dmrt1*Δ13^+/–^ double heterozygous fishes, which were intercrossed to obtain double homozygous mutants used in this study.

### Immunohistochemistry (IHC)

Adult testes were isolated and fixed with 4% PFA (pre-warmed at room temperature (RT)) for 2 hrs at RT. Embryos and larvae were fixed with 4% pre-warmed PFA for 1 hr at RT. Before fixation, eggs were dechorionated with hatching enzyme purchased from NBRP Medaka (https://shigen.nig.ac.jp/medaka/top/top.jsp), and larval skin at the ventral side was opened by forceps. Fixed samples were washed five times in PTW (PBS with 0.1% Tween-20) for 5 min at RT. For larval samples, the skin around gonadal region was removed by forceps and head was cut for genomic lysate preparation for genotyping. For immunostaining, the samples were blocked in 20% Blocking One (Nacalai tesque) in PTW for 2 hrs at RT, and treated with primary antibodies overnight at 4°C. After washing three times in PTW for 30 min at 4°C, the samples were treated with secondary antibodies and DAPI overnight at 4°C. Immunostained samples were washed with PTW for 10 min at RT and mounted on slides with Vectashield (Vector laboratories). Fluorescent images were acquired using Olympus FV1000 confocal scanning microscope. Volocity imaging software was used for 3D reconstruction of confocal images.

Primary antibodies for the following proteins were used: medaka VASA (Aoki et al., 2008) (1:100; rat antiserum, lab-made), α-tubulin (1:100; mouse, SIGMA, T6557), acetylated α-tubulin (1:100; mouse, SIGMA, T6793), γ-tubulin (1:100, rabbit, GeneTex, GTX113286). The secondary antibodies: Alexa Fluor 488-, 568-, and 647-conjugated goat antibodies (1:200, Thermo Fisher Scientific).

### In situ hybridization (ISH)

cDNAs of *buc* and *bucl* were amplified from ovarian cDNA library by PCR using T7-promoter-tagged primers (**Table S1**). After gel-purification, the PCR products were used as templates for *in vitro* transcription of DIG-labeled RNA probe using T7 RNA polymerase (SIGMA) and DIG RNA Labeling Mix (SIGMA) according to the manufacturer’s instructions.

Adult ovaries were fixed with 4% PFA and dehydrated in 100% methanol as described in IHC section. The samples were sequentially washed in 75%, 50%, and 25% methanol in PTW, and twice in PTW at RT for 5 min in each. Then the samples were treated with 10 µg/mL Proteinase K (Roche) in PTW for 15 min at RT, then fixed with 4% PFA for 20 min at RT. After washing five times with PTW for 5 min, the samples were pre-hybridized in hybridization buffer (50% formamide containing 5x SSC, 0.1% Tween-20, 5 mg/mL torula RNA (SIGMA), and 100 µg/mL heparin sulfate) for 2 hrs at 65°C, followed by hybridization with heat-denatured DIG-labeled RNA probe diluted in hybridization buffer at 65°C overnight. Hybridized samples were washed twice with 2x SSCT in 50% formamide for 30 min, twice in 2x SSCT for 15 min, and twice in 0.2x SSCT for 30 min at 65°C. Then the samples were blocked with 5% sheep serum in PTW for 2 hrs at RT. The buffer was replaced by anti-Digoxigenin-AP Fab fragments (1:200, Merk, 11093274910) in 5% sheep serum / PTW overnight at 4°C. After washing six times with PTW for 10 min and twice in staining buffer (100mM Tris-HCl (pH 9.5), 100 mM NaCl, 50mM MgCl_2_, 0.1% Tween-20) for 5 min, signal was detected by 2% NBT/BCIP (Merk, 11681451001) in staining buffer. Signals were observed by KEYENCE BZ-X microscope.

### Fluorescent ISH (FISH)–IHC

FISH-IHC follows ISH protocol up to the blocking step. The samples were treated with antibodies for acetylated α-tubulin (1:100; mouse, SIGMA, T6793), medaka VASA (Aoki et al., 2008) (1:100; rabbit antiserum, lab-made), and Digoxigenin-POD (1:100; Boehringer, 1207733), in 5% sheep serum at 4°C overnight. After washing three times with PTW, anti-DIG-POD signal was amplified with TSA Biotin System Kit (Perkin Elmer) according to the manufacturer’s instruction. Then the samples were treated with secondary antibodies (anti-mouse IgG-Alexa 647 (1:100), anti-rabbit IgG-Alexa 488 (1:100), streptavidin-Alexa 568 (1:100)) and DAPI in 5% sheep serum at 4°C overnight. Fluorescent signals were detected by Olympus FV1000 confocal scanning microscope.

### Quantifying germ cell number and statistical analysis

For quantification of germ cells with the MT dome, VASA-positive cells in whole gonads were counted. For 10 dph XX gonads, meiotic oocytes with condensed chromosomes and small nucleoli were excluded from quantification. Four to six embryos per different developmental stages were counted for quantification. Data are presented as mean ± SD. Data were analyzed by a two-sided paired Welch’s *t*-test.

### Transmission electron microscopy

For TEM analysis, 10 dph XX larvae were dissected and fixed with 0.1M cacodylic acid buffer containing 2% PFA, 2% glutaraldehyde and 0.1% picric acid. Sample preparation and image acquisition were performed in Tokai Electron Microscopy, Inc.

## Acknowledgements

We thank NBRP Medaka for providing a transgenic medaka “Pbeta-actin-tagCFP-mito” (strain ID: TG1020) and hatching enzyme. We are grateful to all members of Tanaka’s laboratory for helpful discussions and suggestions in the study.

---

**Supplementary figure 1.** Immunofluorescence of 0 dph XX (A, C) and 0 dph XY (B, D) gonads stained for γ-tubulin and acetylated α-tubulin (A, B) or pan α-tubulin (C, D). Scale bars: 30 μm (A, B) or 10 μm (C, D). Arrowheads indicate the MT domes formed around γ-tubulin-positive centrioles.

**Supplementary figure 2.** (A) Three-dimensional reconstruction of a 10 dph XY germ cell (dotted line) stained with antibodies against acetylated α-tubulin (green) and γ- tubulin (red). Centrioles are located at the base of the MT dome. (B) Horizontal and vertical sections of the 3D image are shown in A. The MT dome has a hollow inside.

**Supplementary figure 3.** Immunohistochemistry of embryos at stage 28 (A; 30-somite stage), stage 30 (B; 35-somite stage), stage 33 (C; notochord vacuolization stage), and stage 35 (D; visceral blood vessels formation stage) stained for pan α-tubulin, γ-tubulin, and VASA. At stage 30, PGCs come in touch with gonadal somatic precursors, and move dorsally to the prospective gonadal region by stage 33. Expression of the sex-determining gene, *DMY*/*dmrt1bY*, at stage 33 triggers sexual differentiation of gonads by stage 35. Arrowheads indicate the MT domes in the PGCs (A, B) or gonocytes (C, D). Scale bars: 10 μm.

**Supplementary figure 4.** Immunohistochemistry of 10 dph XX (A) and XY (B) gonads. All mitotic germ cells possess the MT dome (arrowheads). Magnified images of mitotic germ cells are shown in A’ and B’. Scale bars: 30 μm (A, B) or 10 μm (A’, B’).

**Supplementary figure 5.** TEM images of type-I oogonia (A, B) and a zygotene oocyte (C). Germ cells are colored pink. Arrowheads indicate Golgi apparatus disposed in a circular manner. Arrows indicate germinal vesicles. Golgi’s circular disposition and germinal vesicles are magnified in A’-C’ and A″-C″, respectively. Arrows in A″-C″ indicate germinal vesicles. Asterisk in C’ indicates a centriole. Scale bars: 2 μm (A-C) or 1 μm (A’-C’, A″-C″).

**Supplementary figure 6.** Genomic loci of *bucky ball* (*buc*) and its paralogs *bucky ball-like* (*bucl*). Predicted gene organization based on Ensembl genome browser release 109 (black; Cuunningham *et al*., 2022) and on our RNA-seq read mapping (red; Kikuchi *et al*., 2019) are indicated. RNA-seq read coverages of medaka XX type-I/II germ cells (three replicates) for *buc* and *bucl* are shown at the bottom.

**Supplementary figure 7.** Syntenic map of *buc* and its paralogs *bucl* and *buc2l* (black, blunt end: 5’). Species and chromosomes are indicated on the left. Gray and red lines connect orthologs between species. Synteny of gray genes is conserved from Spotted gar.

**Supplementary figure 8.** *In situ* hybridization of Bb components, *buc* (A-D) and *bucl* (E-H). NBT/BCIP signals are shown in blue. Expression patterns are summarized in I. Expression of both genes was detected in meiotic oocytes. Transcripts of *buc* and *butl* were localized to Bb (arrowheads) in previtellogenic oocytes. OG: oogonia, OCl: leptotene oocyte, OCz: zygotene oocyte, OCp: pachytene oocyte, OCd: diplotene oocyte, OCpv: previtellogenic oocyte. Developmental stages of germ cells were determined as follows. OG: nucleolus (faintly stained with neutral red) is located at the center of nucleus. OCl: nucleolus is located at nuclear periphery. OCz: nucleolus is located at nuclear periphery, and condensed chromosomes are visible in nucleus. OCp: condensed chromosomes are still visible in nucleus, and cellular size increases. OCd: chromosome decondense, and cellular and nuclear size increase up to 90 µm in diameter. OCpv: cellular size increases up to 150 µm in diameter. Scale bars: 10 μm.

**Movie 1**. 3D reconstruction of acetylated α-tubulin signal in stem-type spermatogonia. The MT dome is constructed at the perinuclear cytoplasmic region. The same cell is shown in Figure S4B’.

**Movie 2**. 3D reconstruction of acetylated α-tubulin signal in a cyst of differentiating oogonia. The MT domes are connected to each other through intercellular bridges. The same cyst is shown in Figure 1B.

**Movie 3**. 3D reconstruction of acetylated α-tubulin signal in a cyst of leptotene oocytes. Microtubules connecting the MT domes were thinner than those in differentiating oogonia. The same cyst is shown in Figure 1C.

**Movie 4**. 3D reconstruction of acetylated α-tubulin signal in a cyst of zygotene oocytes. The MT domes were no longer connected at this stage. The same cyst is shown in Figure 1D.

**Movie 5**. 3D reconstruction of acetylated α-tubulin signal in pachytene oocytes. A cage-like microtubule structure was formed around the nucleus. The same cells are shown in Figure 1E.

**Movie 6**. 3D reconstruction of acetylated α-tubulin signal in diplotene oocytes. A cage-like structure is maintained at this stage.

## References

Abby, E., Tourpin, S., Ribeiro, J., Daniel, K., Messiaen, S., Moison, D., Guerquin, J., Gaillard, J.-C., Armengaud, J., Langa, F., et al. (2016). Implementation of meiosis prophase I programme requires a conserved retinoid-independent stabilizer of meiotic transcripts. Nat. Commun. 7, 10324.

Aoki, Y., Nagao, I., Saito, D., Ebe, Y., Kinjo, M. and Tanaka, M. (2008). Temporal and spatial localization of three germline-specific proteins in medaka. Dev. Dyn. 237, 800–807.

Bontems, F., Stein, A., Marlow, F., Lyautey, J., Gupta, T., Mullins, M. C. and Dosch, R. (2009). Bucky ball organizes germ plasm assembly in zebrafish. Curr. Biol. 19, 414–422.

Chikashige, Y., Tsutsumi, C., Yamane, M., Okamasa, K., Haraguchi, T. and Hiraoka, Y. (2006). Meiotic proteins bqt1 and bqt2 tether telomeres to form the bouquet arrangement of chromosomes. Cell. 125, 59–69.

Cox, R. T. and Spradling, A. C. (2003). A Balbiani body and the fusome mediate mitochondrial inheritance during *Drosophila* oogenesis. Development. 130, 1579–1590.

Cunningham, F., Allen, J. E., Allen, J., Alvarez-Jarreta, J., Amode, M. R., Armean, I. M., Austine-Orimoloye, O., Azov, A. G., Barnes, I., Bennett, R., et al. (2022). Ensembl 2022. Nucleic Acids Res. 50, D988–D995.

Elkouby, Y. M., Jamieson-Lucy, A. and Mullins, M. C. (2016). Oocyte polarization is coupled to the chromosomal bouquet, a conserved polarized nuclear configuration in meiosis. PLoS. Biol. 14, e1002335.

Elkouby, Y. M. (2017). All in one - integrating cell polarity, meiosis, mitosis and mechanical forces in early oocyte differentiation in vertebrates. Int. J. Dev. Biol. 61(3-4-5), 179–193.

Ge, X., Grotjahn, D., Welch, E., Lyman-Gingerich, J., Holguin, C., Dimitrova, E., Abrams, E. W., Gupta, T., Marlow, F. L., Yabe, T., et al. (2014). Hecate/Grip2a acts to reorganize the cytoskeleton in the symmetry-breaking event of embryonic axis induction. PLoS. Genet. 10, e1004422.

Gill, M. E., Hu, Y.-C., Lin, Y. and Page, D. C. (2011). Licensing of gametogenesis, dependent on RNA binding protein DAZL, as a gateway to sexual differentiation of fetal germ cells. Proc. Natl. Acad. Sci. U. S. A. 108, 7443–7448.

Iwamatsu, I. (2004). Stages of normal developmental in the medaka *Oryzias latipes*. Mech. Dev. 121, 605–618.

Jamieson-Lucy, A. and Mullins, M. C. (2019). The vertebrate Balbiani body, germ plasm, and oocyte polarity. Curr. Top. Dev. Biol. 135, 1–34.

Kikuchi, M., Nishimura, T., Saito, D., Shigenobu, S., Takada, R., Gutierrez-Triana, J. A., Cerdán, J. L. M., Takada, S., Wittbrodt, J., Suyama, M. and Tanaka, M. (2019). Novel components of germline sex determination acting downstream of *foxl3* in medaka. Dev. Biol. 445, 80–89.

Kikuchi, M., Nishimura, T., Ishishita, S., Matsuda, Y. and Tanaka, M. (2020). *foxl3*, a sexual switch in germ cells, initiates two independent molecular pathways for commitment to oogenesis in medaka. Proc. Natl. Acad. Sci. U. S. A. 117, 12174–12181.

Kikuchi, M. and Tanaka, M. (2022). Functional modules in gametogenesis. Front. Cell. Dev. Biol. 10, 914570.

Kurokawa, H., Aoki, Y., Nakamura, S, Ebe, Y., Kobayashi, D. and Tanaka, M. (2006). Time-lapse analysis reveals different modes of primordial germ cell migration in the medaka *Oryzias latipes*. Dev. Growth. Differ. 48, 209–221.

Lei, L. and Spradling, A. C. (2016). Mouse oocytes differentiate through organelle enrichment from sister cyst germ cells. Science. 352, 95–99.

Lu F.-I., Thisse, C. and Thisse, B. (2011). Identification and mechanism of regulation of the zebrafish dorsal determinant. Proc. Natl. Acad. Sci. U. S. A. 108, 15876–15880.

Marlow, F. L. and Mullins, M. C. (2008). Bucky ball functions in Balbiani body assembly and animal-vegetal polarity in the oocyte and follicle cell layer in zebrafish. Dev. Biol. 321, 40–50.

Masuyama, H., Yamada, M., Kamei, Y., Fujiwara-Ishikawa, T., Todo, T., Nagahama, Y. and Matsuda, M. (2012). *Dmrt1* mutation causes a male-to-female sex reversal after the sex determination by *Dmy* in the medaka. Chromosome. Res. 20, 163–176.

Matsuda, M., Nagahama, Y., Shinomiya, A., Sato, T., Matsuda, C., Kobayashi, T., Morrey, C. E., Shibata, N., Asakawa, S., Shimizu, N., et al. (2002). DMY is a Y-specific DM-domain gene required for male development in the medaka fish. Nature. 417, 559–563.

Nakamura, S., Kobayashi, D., Aoki, Y., Yokoi, H., Ebe, Y., Wittbrodt, J. and Tanaka, M. (2006). Identification and lineage tracing of two populations of somatic gonadal precursors in medaka embryos. Dev. Biol. 295, 678–688.

Nakamura, S., Kobayashi, K., Nishimura, T., Higashijima, S. and Tanaka, M. (2010). Identification of germline stem cells in the ovary of the teleost medaka. Science. 328, 1561–1563.

Nanda, I., Kondo, M., Hornung, U., Asakawa, S., Winkler, C., Shimizu, A., Shan, Z., Haaf, T., Shimizu, N., Shima, A., Schmid, M. and Schartl, M. (2002). A duplicated copy of DMRT1 in the sex-determining region of the Y chromosome of the medaka, *Oryzias latipes*. Proc. Natl. Acad. Sci. U. S. A. 99, 11778–11783.

Nishimura, T. and Tanaka, M. (2014). Gonadal development in fish. Sex. Dev. 8, 252–261.

Nishimura, T., Sato, T., Yamamoto, Y., Watakabe, I., Ohkawa, Y., Suyama, M., Kobayashi, S. and Tanaka, M. (2015). *foxl3* is a germ cell-intrinsic factor involved in sperm-egg fate decision in medaka. Science. 349, 328–331.

Nishimura, T. and Tanaka, M. (2016). The mechanism of germline sex determination in vertebrates. Biol. Reprod. 95, 30.

Nishimura, T., Yamada, K., Fujimori, C., Kikuchi, M., Kawasaki, T., Siegfried, K. R., Sakai, N. and Tanaka, M. (2018). Germ cells in the teleost fish medaka have an inherent feminizing effect. PLoS. Genet. 14, e1007259.

Niu, W. and Spradling, A. C. (2022). Mouse oocytes develop in cysts with the help of nurse cells. Cell. 185, 2576–2590.e12.

Nojima, H., Rothhämel, S., Shimizu, T., Kim, C.-H., Yonemura, S., Marlow, F. L. and Hibi, M. (2010). Syntabulin, a motor protein linker, controls dorsal determination. Development. 137, 923–933.

Pepling, M. E., Wilhelm, J. E., O’Hara, A. L., Gephardt, G. W. and Spradling, A. C. (2007). Mouse oocytes within germ cell cysts and primordial follicles contain a Balbiani body. Proc. Natl. Acad. Sci. U. S. A. 104, 187–192.

Roper, K. and Brown, N. H. (2004). A spectraplakin is enriched on the fusome and organizes microtubules during oocyte specification in *Drosophila*. Curr. Biol. 14, 99–110.

Saito, D., Morinaga, C., Aoki, Y., Nakamura, S., Mitani, H., Furutani-Seki, M., Kondoh, H. and Tanaka, M. (2007). Proliferation of germ cells during gonadal sex differentiation in medaka: Insights from germ cell-depleted mutant zenzai. Dev. Biol. 310, 280–290.

Sakuma, T., Hosoi, S., Woltjen, K., Suzuki, K.-I., Kashiwagi, K., Wada, H., Ochiai, H., Miyamoto, T., Kawai, N., Sasakura, Y., et al. (2013). Efficient TALEN construction and evaluation methods for human cell and animal applications. Genes. Cells. 18, 315–326.

Scherthan, H. (2001). A bouquet makes ends meet. Nat. Rev. Mol. Cell. Biol. 2, 621–627.

Shibata, N. and Hamaguchi, S. (1988). Evidence for the sexual bipotentiality of spermatogonia in the fish, *Oryzias latipes*. J. Exp. Zool. 245, 71–77.

Shibuya, H., Morimoto, A. and Watanabe, Y. (2014). The dissection of meiotic chromosome movement in mice using an in vivo electroporation technique. PLoS. Genet. 10, e1004821.

Soh, Y. Q. S., Mikedis, M. M., Kojima, M., Godfrey, A. K., Rooij, D. G. and Page, D. C. (2017). *MeioC* maintains an extended meiotic prophase I in mice. PLoS. Genet. 13, e1006704.

Sumita, R., Nishimura, T. and Tanaka, M. (2022). Dynamics of spermatogenesis and change in testicular morphology under ’mating’ and ’non-mating’ conditions in medaka (*Oryzias latipes*). Zoolog. Sci. 38, 436–443.

